# Genomic Characterisation of Group B *Streptococcus* from Argentina: Insights into Prophage Diversity, Virulence Factors and Antibiotic Resistance Genes

**DOI:** 10.1101/2024.12.20.629649

**Authors:** Veronica Kovacec, Sabrina Di Gregorio, Mario Pajon, Uzma Basit Khan, Tomas Poklepovich, Josefina Campos, Chiara Crestani, Stephen D. Bentley, Dorota Jamrozy, Marta Mollerach, Laura Bonofiglio

## Abstract

Group B *Streptococcus* (GBS) is a commensal bacterium that can cause severe infections in infants and adults with comorbidities. Resistance and reduced susceptibility to antibiotics is continually on the rise, and vaccines remain in-development. Prophages have been reported to contribute to GBS evolution and pathogenicity. However, no studies are available to date on prophage contribution to the epidemiology of GBS isolates from humans in South America. In the context of an Argentinian multicentric study, we had previously phenotypically characterised 365 human GBS isolates from invasive disease, urinary infections and maternal colonisation. These isolates had been whole genome sequenced and their prophage presence bioinformatically determined. In this study, we genomically characterised the isolates and analysed the prophage content in the context of the epidemiological data. The phylogenetic analysis of the 365 genomes with 103 GBS from public databases revealed that Argentinian GBS were related to isolates from around the world. The most prevalent lineages, independently of the isolate source, were CC23/Ia and CC12/Ib. Genes encoding virulence factors involved in immune response evasion, tissue damage and adherence to host tissues and invasion were found in all of the genomes in accordance with previously described lineage distribution. According to the prevalent capsular types and the distribution of specific virulence factors in Argentinian GBS, over 95% coverage would be expected from the vaccines currently under development. Antibiotic resistance determinants (ARDs) to at least one antibiotic class were found in 90% of the genomes, including novel mutations in *pbp*2x, while more than 15% carried ARDs to 3 or more classes. GBS collected from urinary infections carried a significantly higher proportion of ARDs to multiple antibiotic classes than the rest of the isolates. A total of 454 prophages were found among the 468 genomes analysed, which were classified into 23 prophage types. Prophage presence exhibited variations based on GBS clonal complex and capsular type. A possible association between an increased GBS pathogenicity and the carriage of prophages with integrase type GBS*Int*8 and/or the presence of genes that encode the Phox Homology domain has been observed. The highest prevalence of prophages per genome was found in lineages CC17/III and CC19/III, while the lowest amount was observed in CC12/Ib. Overall, the highest density of prophages, virulence factors and ARDs determinants was found in CC19 isolates, mostly of capsular type III, independently of the isolates source. This is the first analysis of the human-associated GBS population in South America based on whole genome sequencing data, which will make a significant contribution to future studies on the global GBS population structure.

**Data summary:** Supplementary results and figures can be found in Supplementary Material 1. Supplementary tables can be found in Supplementary Material 2. All datasets analysed in this study are detailed in the Supplementary Materials. Metadata about the genomes analysed can be found in the microreact project created for this study: https://microreact.org/project/gbs-pangenomic-analysis.

**Impact statement:** In Latin America studies on the epidemiology of Group B Streptococcus (GBS) are scarce, especially those describing clonal complex and serotype distribution, and the role of prophages in GBS epidemiology has not been studied. This article addresses the first genomic characterisation of the human-isolated GBS population in Latin America based on whole genome sequencing data, with special focus on the analysis of prophage content. We determine the clonal complex and serotype distribution of 365 GBS isolates collected from clinical samples in an Argentinian multicenter study and analyse the presence of prophages and virulence and antibiotic resistance determinants in the context of the epidemiological data. Through these analyses, we were able to determine how GBS population structure in Argentina differs from other parts of the world and to predict the potential coverage of the in-development GBS vaccines. We also found a possible association between the carriage of certain types of prophages and an increased GBS pathogenicity. In the context of increased global efforts to develop new strategies to prevent GBS infections through vaccine development, this study makes a significant contribution to our understanding of the global GBS population structure.

## Introduction

*Streptococcus agalactiae* (group B *Streptococcus*; GBS) is a commensal bacterium that colonises the human intestinal and genitourinary tracts. GBS is the leading cause of neonatal sepsis and other perinatal infections such as meningitis and pneumonia (1). In recent decades, there has been a marked increase in the incidence of invasive infections caused by GBS in non-pregnant individuals, especially in the elderly and those suffering from underlying medical conditions (2–8). Advanced age, diabetes *mellitus* and cancer have been postulated as the main risk factors for GBS invasive disease in adult patients (7,9,10).

Penicillin is the first-line drug for the treatment and prevention of GBS infections and combination of both penicillin and gentamicin is indicated for sepsis in infants (11,12). In patients with penicillin allergy macrolides/lincosamides are used (13,14), while vancomycin is reserved for cases of macrolides/lincosamides resistance (15). Macrolide and lincosamide resistance has been gradually increasing in the last decades (16–20). Resistance to other antibiotic families has also emerged, such as reduced susceptibility to beta-lactams (21) as well as resistance to aminoglycosides (22), vancomycin (23) and fluoroquinolones (24). This has been associated with dissemination of multidrug resistant (MDR) clones (2–5,8,18).

To date, there are no approved vaccines for the prevention of GBS infections, but there are several serotype-specific or protein-based vaccines in development (25). The impact of the vaccine-based prevention will depend on vaccine coverage according to the selected antigens, so it is imperative to study the diversity of GBS isolates circulating in each region. An international consortium called Juno was established to analyse the diversity, vaccine target distribution and genetic determinants associated with GBS infection worldwide (https://www.gbsgen.net/).

Prophages are important vehicles for horizontal gene transfer (HGT) (26) and they play a significant role in bacterial evolution by introducing genes that enhance bacterial fitness and virulence (27,28). Up to 20% of bacterial genomes can be constituted by prophages and it has been described that pathogenic strains tend to carry more phage-related genes than non-pathogenic strains (29–31), which was also observed for GBS (32).

Recent studies on the role of lysogeny in the evolution and pathogenicity of human GBS isolates revealed that acquisition of certain prophages (of possible animal origin) might have been related to the emergence of specific GBS clones that were pathogenic for infants or adults in Europe (32–35). Whether such association between prophage content and GBS pathogenicity is present in other geographical areas is unknown.

We previously analysed the prophage content of 365 GBS genomes from Argentina, detecting 325 prophages, which were classified into 19 prophage types, and found significant associations between the prevalent prophage types and certain GBS clonal complexes (36). The present study aims to characterise further genomic diversity of the 365 Argentinian GBS isolates as well as the prophage presence in the context of the epidemiological data.

## Methods

### Genomes used in this study

Within the framework of a Argentinian national multicentric study that involved 40 health centres in 12 provinces of Argentina and took place in 2014-2015, we collected 450 GBS isolates from invasive or urinary infections as well as colonised pregnant women. All isolates had been previously characterised phenotypically and genotypically by PFGE (3–5) and 365/450 were whole genome sequenced as part of the Juno project. The geographic coverage of the 365 Argentinian GBS genomes can be visualized in the microreact project created for this study (https://microreact.org/project/gbs-pangenomic-analysis). Sequencing and raw sequence data handling methodology were described previously (36). The dataset was divided into four collections. The infant invasive collection (iiGBS, N=22) from infants between 1 and 90 days of age with an invasive infection (13/22 from early-onset disease - EOD; 9/22 from late-onset disease-LOD). The non-infant invasive collection (niGBS, N=133) from children over 3 months old (N=3) and adults (N=130). The urinary infection collection (uGBS, N=145) from adolescents (aged 16 to 17 years, N=4) and adults (N=141). The colonising collection (cGBS, N=85) from adolescent (aged 14 to 17 years, N=8) and adult (N=77) pregnant women.

The dataset was supplemented with 103 whole genome GBS assemblies from humans collected in 16 countries across 5 continents, retrieved from NCBI under the category of RefSeq (https://www.ncbi.nlm.nih.gov/, accessed in August 2022), (Supplementary Table S1). Assemblies were selected based on the following criteria: 1) isolates from children or adults suffering from invasive or urinary infections, or from colonisation; 2) isolates representing clonal complexes common among GBS from infections in humans in Argentina (CC23, CC17, CC19, CC1, CC12, CC452, data from this work); 3) isolates collected during time period that aligned with the sampling period of our isolates. For more information about these isolates see the <u>microreact project</u>.

### GBS genomic characterisation

The steps followed for GBS characterisation are detailed in this section and summarised in Supplementary Figure 1.

To reconstruct a core-SNP phylogeny, the 468 assemblies (365 from Argentina plus 103 from the global dataset) were annotated with Prokka v1.14.5 (37) and a core gene alignment was generated with Roary v3.13.0 (38). Single nucleotide polymorphisms (SNPs) were identified with SNP-sites v2.5.1 (39). A maximum-likelihood (ML) phylogenetic tree was reconstructed using IQ-TREE v1.6.12 (40), with automatic selection of the substitution model and 1000 SH-aLRT (41) and 1000 ultrafast-bootstrap (42) replicates for branch support analysis.

Capsular types were determined by sequence similarity between the assemblies and reference capsular gene sequences of the 10 GBS capsular types, through a blastn (BLAST v2.9.0+, https://blast.ncbi.nlm.nih.gov/Blast.cgi) search (E value threshold of 1e^-100,^ otherwise default parameters), as described by Sheppard (43). As these authors did not publish the reference sequence of the serotype IX capsular gene, the *cps*O gene sequence used by Breeding (44) for capsular type IX detection by PCR was incorporated into the capsular gene database. In all cases, a capsular type was assigned when there was at least 95% of sequence identity over 90% of the sequence length. The results were compared with those previously obtained by phenotypic serotyping conducted on the same isolate collection (3,5,4). A comparative analysis between the capsular type loci (*cps*) of non-typable isolates and reference strains (45) of each capsular type (Supplementary Table S2) was performed with clinker v0.027 (46).

MLST was determined with the software mlst v2.22.1 (47). The pubMLST website (https://pubmlst.org/organisms/streptococcus-agalactiae) (48,49) was used for sequence type (ST) assignment when new alleles were detected and to assign each ST to a clonal complex (CC).

Prophage screening and typing was performed with a methodology previously developed for GBS-prophage detection and their classification in prophage types (36), according to their phylogenetic group and integrase type. Shortly, GBS genomes were searched for genes specific to each prophage type by the following methodology: i) blastn search against the Prophage-group database (36); a positive result was considered when at least one of the prophage group-specific genes was detected with at least 75% of sequence identity over 75% of the sequence length; in the special case of group A prophages, at least two genes should be detected: *hha*I **or** *clp*P **and** the gene coding for the holin **or** the lysin. ii) blastx search against the Prophage-integrase database (36,50); a positive result was considered when the integrase gene was detected with at least 90% of sequence identity over 95% of the sequence length. iii) Prophage classification by prophage type: integration of classifications by prophage group/integrase type.

Virulence determinants (VDs) were searched with Abricate v0.9.9 (51), using the VFDB database v2023-06-27 (52) and by a blastn v2.9.0+ search against a reference database of genes coding for surface proteins involved in GBS virulence (53). In all cases, the threshold for gene detection was established as 80% identity over 80% of the reference sequence length.

Antibiotic resistance determinants (ARDs) were searched with Abricate v0.9.9, using the ResFinder database v2023-06-27 (54) and AMRFinderPlus v3.11.4, database v2023-04-17.1 (55). Additionally, a blastn v2.9.0+ search was performed against two reference databases created by Metcalf (56) for specific detection of ARDs in GBS genomes, including known variants of the PBP2x transpeptidases (57) and resistance determinants to quinolones and several other antibiotic families (58). In all cases, the threshold for gene detection was established as 80% identity over 80% of the reference sequence length. In the case of allelic variants of genes coding for PBP2x or the genes involved in quinolone resistance (*par*C and *gyr*A), novel variants were considered those without 100% identity with any reference variant. The results were compared with those previously obtained by phenotypic analysis (3,5,59).

### Integration of information

The Microreact application (60) was used for an integral visualisation of the genomic analysis results. All the information collected was evaluated in the context of both the whole set of genomes and each of the Argentinian GBS collections.

### Statistical analysis

Chi-squared and Fisher’s exact test (two-tailed) were used to evaluate distribution and correlation of categorical variables. A *p*-value of ≤0.05 was considered to be significant.

## Results

### Phylogenetic and lineage analysis of GBS genomes

The core-SNP phylogeny showed that Argentinian isolates clustered with the globally derived human GBS. Cluster distribution correlated with CC and, in most cases, with capsular type (Figure 1).

**Figure 1.**
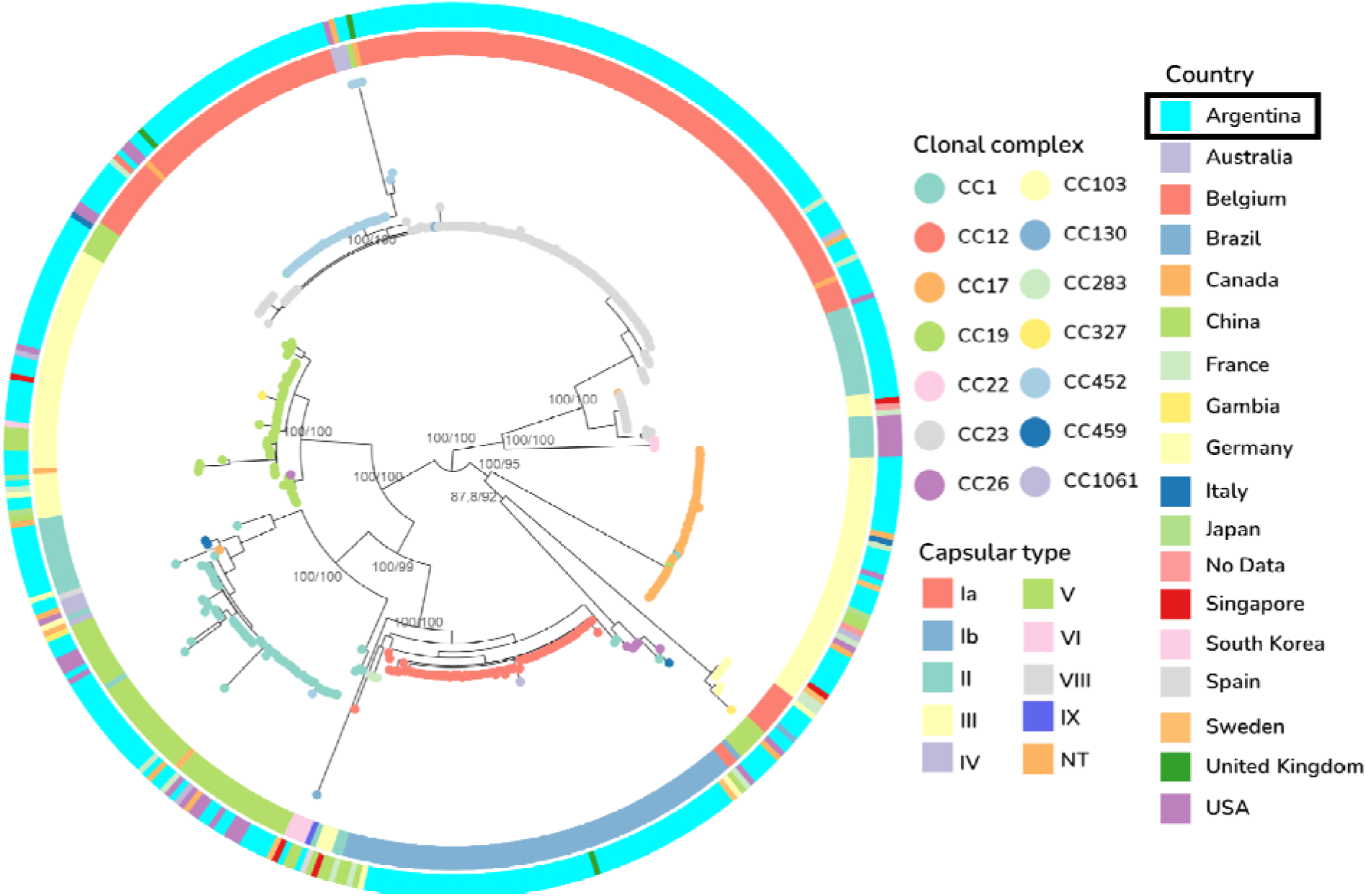
Maximum likelihood phylogenetic tree of the 468 GBS genomes analysed here. The tree was midpoint rooted, with nodes coloured according to clonal complex (CC). The inner ring represents the capsular type and the outer ring the country of isolation. Support values (SH-aLRT/ultrafast Bootstrap) are shown as labels at the roo of the main clades. The dataset is available at: https://microreact.org/project/gbs-pangenomic-analysis

Significant associations (*p*<0.05) were found between the most prevalent capsular types and clonal complexes: CC23/Ia, CC452/Ia, CC12/Ib, CC19/II, CC23/II, CC17/III, CC19/III and CC1/V. Lineages CC23/Ia and CC12/Ib were the most prevalent among all Argentinian isolates (31% and 17%, respectively) and in each of the Argentinian collections (Figure 2). A similar distribution of lineages was observed in each collection, with the exception of the lineage CC19/III, whose frequency was significantly higher in uGBS and lower in niGBS (*p*<0.05 in both cases). Eleven new sequence types (ST) belonging to prevalent CCs were found among Argentinian genomes and submitted to the pubMLST database.

**Figure 2.**
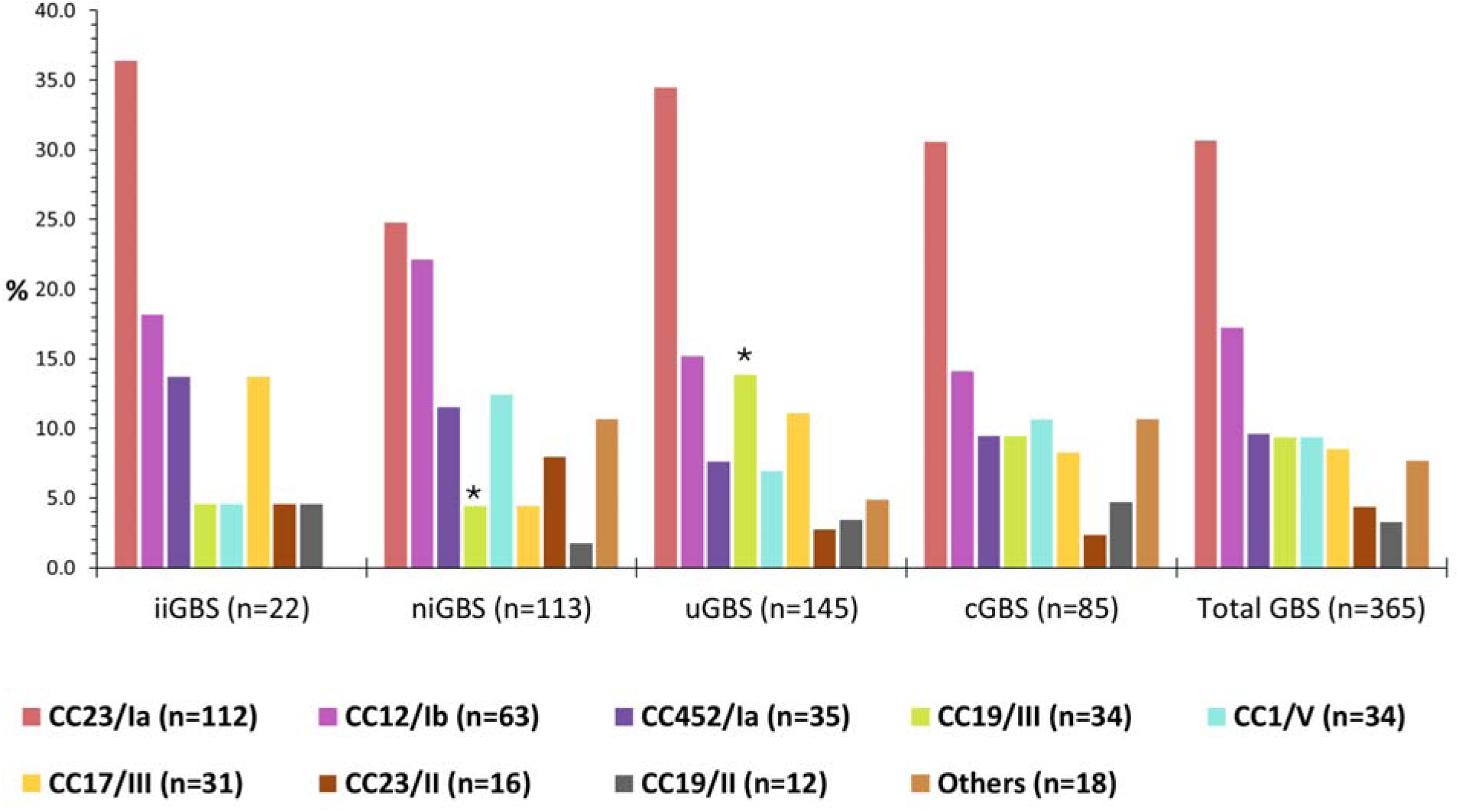
Lineage distribution among Argentinian GBS isolates according to collection. iiGBS: GBS recovered from infant invasive infections; niGBS: GBS recovered from non-infant invasive infections; uGBS: GBS recovered from urinary infections; cGBS: GBS recovered from pregnant women during prenatal screenings. The lineages that differ significantly (*p*<0.05) between collections are marked with *.

### Analysis of capsular type discrepancies and non-typeable GBS isolates

The comparison between the *in silico* and phenotypic capsular typing of the Argentinian isolates revealed 85% correlation. In the discrepant 15%, there was no correlation between the capsular type and the differing serotype, that is, the same discrepancies were not always found between both methods (Supplementary Table S3). All results shown in this study regarding capsular type were obtained *in silico*, as described in the Methods section.

Five out of the 468 isolates were classified as non-typeable (NT) by *in silico* analysis (Supplementary Table S4). This included three Argentinian GBS, two of which were phenotypically non-typable and one typed as serotype Ia. All NT genomes were searched manually for the *cps* operon (constituted by *cps* and *neu* genes). In 3/5 isolates the failure of capsular typing could be explained by deletion of all *cps* genes, with presence of only *neu* genes. The two remaining NT isolates presented seemingly complete *cps* operons, and were compared against reference sequences of each capsular type (Supplementary Figure 2). Isolate JN_AR_GBS348 (CC19, from Argentina), shared the highest homology (>94%) with capsular type III, but contained a truncated copy of the *cps*J gene (Supplementary Figure 3A). The strain MIN-180 (CC452, from the United Kingdom), shared 100% homology with all capsular type V operon genes, except for the *cps*O gene due to insertion of IS5-like transposases (Supplementary Figure 3B).

### Prophage presence in the context of GBS epidemiology

We reported previously the detection and classification of 325 prophages within this dataset of 365 Argentinian GBS (36). In this study, the prophage presence was analysed in the context of GBS epidemiological data.

A total of 454 prophages were found among the 468 genomes analysed here, and classified into 23 prophage types. The presence of 1 or 2 prophages per isolate was the most common, though some isolates carried up to four prophages. Prophage type distribution was analysed in the context of GBS phylogeny as well as CC and capsular type assignment (Figure 3).

**Figure 3.**
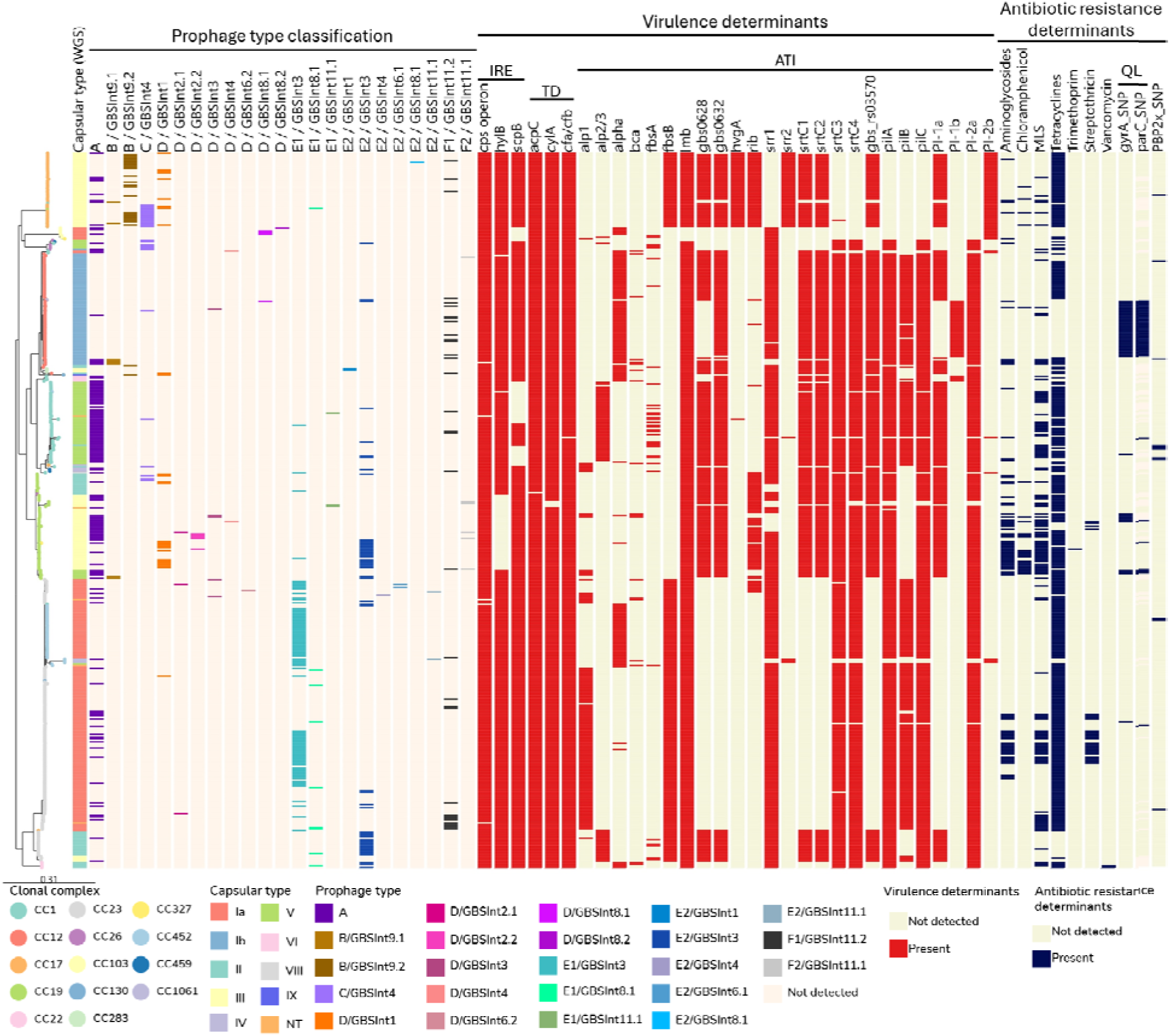
Phylogenetic tree of GBS isolates annotated with the distribution of prophages, virulence determinants and antibiotic resistance determinants. Core-SNP maximum likelihood phylogenetic tree, midpoint rooted, with nodes coloured by clonal complex. Strains metadata are shown as coloured blocks. IRE: Immune response evasion; TD: tissue damage; ATI: adherence to host tissues and invasion; MLS: macrolides/lincosamides/streptogramins; QL: quinolones; NT: non-typeable. The virulence genes of the *cyl* operon, besides *cyl*A and *acp*C, were present in all the genomes and are not included in the figure. https://microreact.org/project/gbs-pangenomic-analysis

Within the predominant lineages, the highest number of prophages per genome was found in CC17/III and CC19/III, while the lowest in CC12/Ib (Figure 3). A significant association was observed (*p*<0.05) between the following lineages and prophage types (Supplementary Figure 4): CC23/Ia with E1/GBS*Int*3 and A; CC12/Ib with F1/GBS*Int*11.2; CC1/V with A; CC19/III with E2/GBS*Int*3, A and D/GBS*Int*1; CC17/III with C/GBS*Int*4, D/GBS*Int*1 and B/GBS*Int*9.2; CC452/Ia with E1/GBS*Int*3; CC23/II with E2/GBS*Int*3.

Interestingly, 13/15 prophages with integrase type GBS*Int*8.1 or GBS*Int*8.2 were detected in isolates from invasive or urinary infections, but were absent in isolates from vaginal carriage. The remaining 2/15 were found in global isolates from unknown origin. These prophages were found in GBS from six different lineages: 4/15 CC103/Ia, 1/15 CC12/Ib, 2/15 CC17/III, 1/15 CC22/II, 6/15 CC23/Ia and 1/15 CC23/II.

While Argentinian isolates from all collections carried predominantly E1/GBS*Int*3, E2/GBS*Int*3 and A prophages, we found significant (p<0.05) associations between cGBS and C/GBS*Int*4 prophages, and between GBS from disease and prophages with integrase type GBS*Int*8.1 (Figure 4). There were no significant differences in the prevalence of genomes lacking a prophage sequence, although a lower prophage content was observed in iiGBS (Supplementary Figure 5).

**Figure 4.**
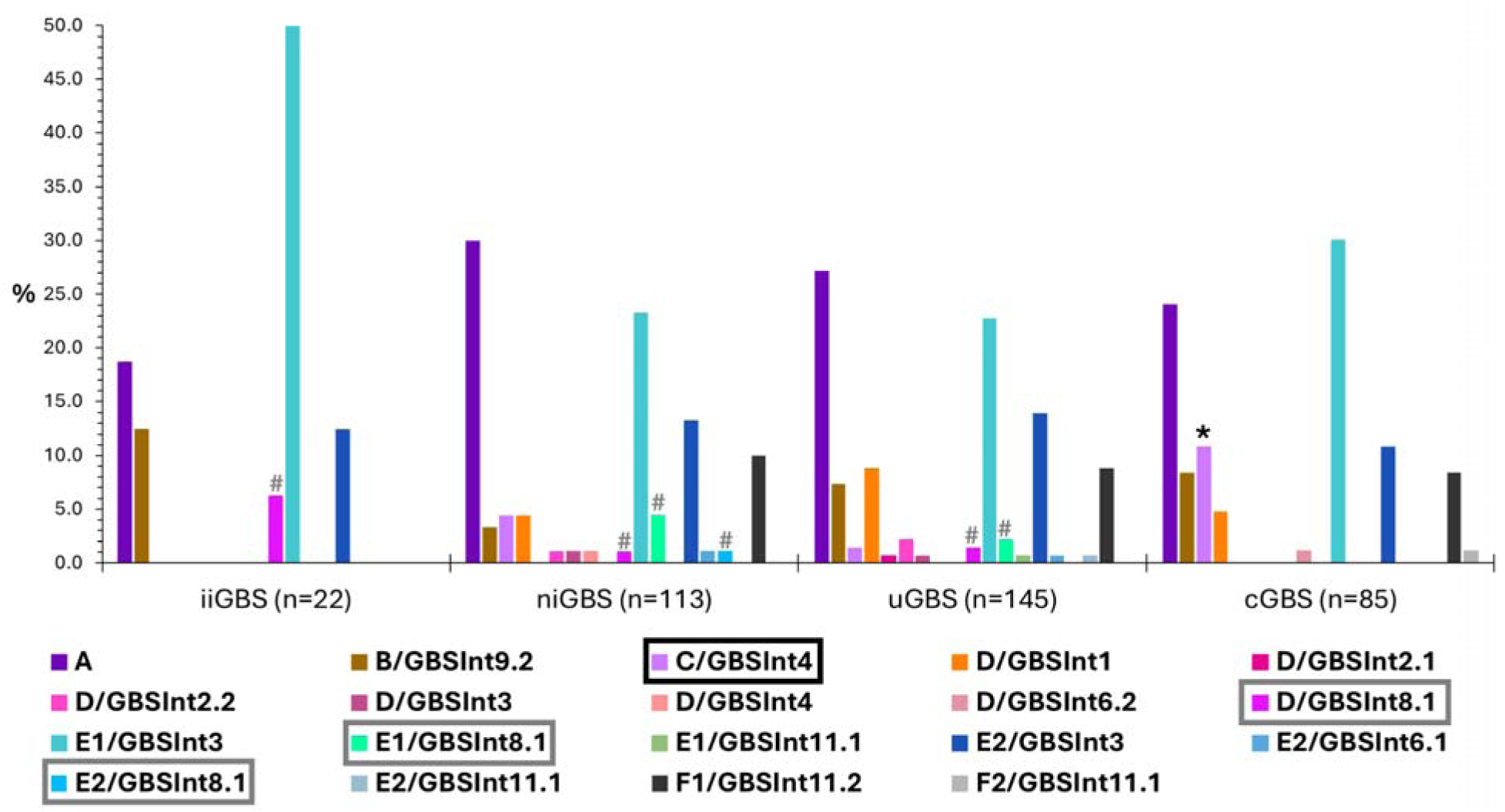
Distribution of prophages in Argentinian GBS isolates according to collection. iiGBS: GBS recovered from infant invasive infections; niGBS: GBS recovered from non-infant invasive infections; uGBS: GBS recovered from urinary infections; cGBS: GBS recovered from pregnant women during prenatal screenings. Significant difference (*p*<0.05) in distribution of a prophage type is shown with an *, and of an integrase type with an #.

Our previous phylogenetic analysis of GBS prophages (36) revealed that 32/87 of type A prophages from Argentinian isolates were related to the type A prophages 12/111phiA and phiStag1, associated with CC17 virulence in infants (33,34,61). In this study we found that these prophages were rare in Argentinian CC17 (1/31, from a uGBS), most (20/32) were carried by 20/112 CC23/Ia isolates and only 1/32 belonged to a iiGBS.

### Presence of VDs and ARDs in the context of GBS epidemiology

We detected the presence of diverse known GBS VDs in all the isolates (n=468), including those involved in immune response evasion (IRE-VD), tissue damage (TD-VD) and adherence to host tissues and invasion (ATI-VD) (Figure 3, for more details see Supplementary Figure 6). The results were in accordance with previous reports on VDs prevalence according to GBS lineage (see ‘‘Supplementary Results’’ in Supplementary Material 1).

Some genes presented different distributions when analysing VD presence according to Argentinian GBS collection. The *hyl*B gene had significantly (*p*<0.001) lower prevalence in niGBS (4%) than in the other collections (over 84%), in accordance with the lower prevalence of lineage CC19/III in niGBS. The *rib* gene prevalence was significantly (*p*<0.01) lower in niGBS (12%, against over 22% in the other collections), which was probably linked to the lower prevalence of serotype III isolates in that collection. The *srt*C3 gene was found in a significantly (*p*<0.01) higher proportion in isolates from invasive infections (92%) compared to those from urinary infections or colonisation (56% and 53%, respectively), and the pilus island PI-2b presented 15% prevalence in invasive GBS versus 32% and 28% in urinary or colonisation isolates, respectively. In the cases of *srt*C3 and PI-2b, no correlation was found between VD distribution and lineage prevalence in each collection.

ARDs to at least one antibiotic class were found in 90% (420/468) of the genomes (Figure 3, for the detailed genes of each family see Supplementary Figure 6). All isolates presented the *mre*A gene (Macrolide efflux pump) (62), independently from Macrolides susceptibility, as previously described (63,64), so *mre*A was not considered an ARD in these results.

None of the VDs and ARDs detected based on gene databases used in this study were found in a prophage.

### ARDs to beta-lactam class

The majority of genomes (457/468) carried common PBP2x variants (1, 2, 4 and 5), not associated with reduced beta-lactam susceptibility (RBLS) (65). Variant 68 of PBP2x, which was previously reported as potentially associated with RBLS (65), was found in 2 Argentinian CC1/V uGBS isolates that were phenotypically susceptible to penicillin. PBP2x variants not previously described were found in 8/468 genomes, all from Argentinian GBS isolates susceptible to penicillin (Table 1). In one GBS genome from China, a PBP2x with a premature stop codon was found, but no data was available about beta-lactam susceptibility (Table 1).

**Table 1.**
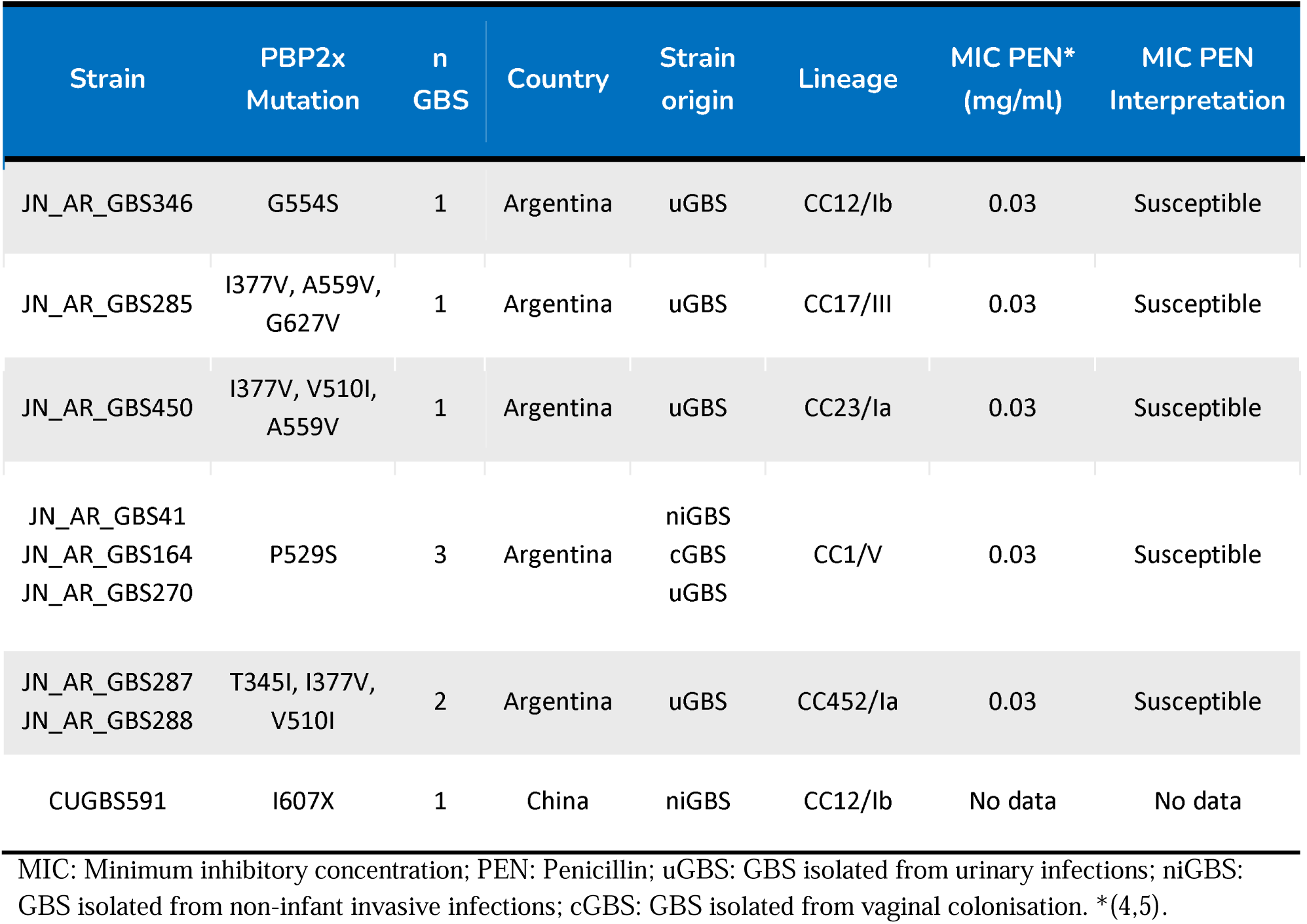
Novel variants of PBP2x found in analysed GBS genomes.

Among Argentinian GBS, the uGBS isolates had a significantly (*p*<0.01) higher prevalence of rare PBP-2x variants in comparison to other collections.

### ARDs to macrolide, lincosamide and streptogramin B (MLS) class

MLS resistance genes were found in 28% (131/468) of the genomes, mostly in lineages CC23/Ia, CC19/III and CC1/V (Figure 3, Supplementary Figure 6). In the Argentinian isolates, their prevalence was 25% (92/365) and no significant difference was found in the content of these genes according to the collection category.

The presence of these genes was analysed in the context of their phenotypic susceptibility to MLS in the 365 Argentinian GBS. Concordance between the phenotypic resistance profile and the ARDs content was found in 97% (354/365) of the isolates. The discrepancies found in the remaining 11/365 genomes are detailed in Supplementary Material 1.

### ARDs to aminoglycoside class

ARDs to aminoglycosides were found in 18% (82/468) of the genomes, predominantly in CC19/III and CC23/Ia (Figure 3, Supplementary Figure 6). In the Argentinian genomes, their prevalence was of 20% (71/365), with a significantly higher (*p*<0.01) frequency in uGBS isolates, while niGBS showed a significantly lower prevalence (*p*<0.01).

Phenotypic susceptibility to streptomycin (STR) and gentamicin (GEN) had been tested for 135 sequenced Argentinian invasive GBS isolates (both from infants and non-infants) (4) and concordance with genotypic profiles was found in 99% (133/135) of the isolates. The discrepancies found in the remaining 2/135 genomes are detailed in Supplementary Material 1.

### ARDs to quinolone class

Point mutations in the quinolone resistance-determining regions (QRDR) of genes *gyr*A and/or *par*C were found in 11% (51/468) of the genomes, most belonging to CC12/Ib and CC19/III. In Argentinian GBS the prevalence was 12% (45/365) and no significant difference was found in the presence of these ARDsby collection. The association between fluoroquinolone resistance and the clonal expansion of ST10/Ib GBS in Argentina was reported in our previous work (3).

Concordance was observed between the phenotypic quinolone-susceptibility profile (3,4) and the bioinformatic detection of point mutations in the *gyr*A and/or *par*C genes in 97% (355/365) of Argentinian GBS. The discrepancies found in the remaining 10/365 genomes are detailed in Supplementary Material 1.

### ARDs to other antibiotic classes

More than 77% of the genomes (365/468; 282/365) carried genes that confer resistance to tetracyclines. The frequency of these genes in the prevalent lineages was higher than 80%, with the exception of CC12/Ib (only present in 43% of the isolates) and CC23/II (no carriage of *tet* genes). High prevalence of *tet* genes is to be expected in GBS isolates, which is why tetracyclines are no longer used for human treatment nor are phenotypically tested for resistance (63).

No ARDs to vancomycin were found in the Argentinian isolates, in accordance with the phenotypic susceptibility previously determined for uGBS and cGBS (4,5). ARDs to vancomycin were only found in 2/468 previously described GBS from the USA (66).

Less than 10% of isolates carried ARDs to antibiotic classes not used for treatment of GBS infections in humans, thus not phenotypically tested in the Argentinian isolates: streptothricin (27/468), chloramphenicol (25/468) and trimethoprim (1/468).

### ARDs to multiple antibiotic classes

The presence of ARDs to three or more antibiotic classes was observed in 15% (71/468) of the genomes (Supplementary Figure 7), so they could be considered multi-drug resistant (MDR) isolates. These isolates belonged to CC19/III (30/71), followed by CC23/Ia (23/71), CC12/Ib (5/71) and CC17/III (4/71) (Figure 5). Mutations in PBP2x potentially associated with RBLS and those not previously described were included in this analysis, as the accumulation of such mutations has been described as a risk factor for RBLS (56,67–69).

**Figure 5.**
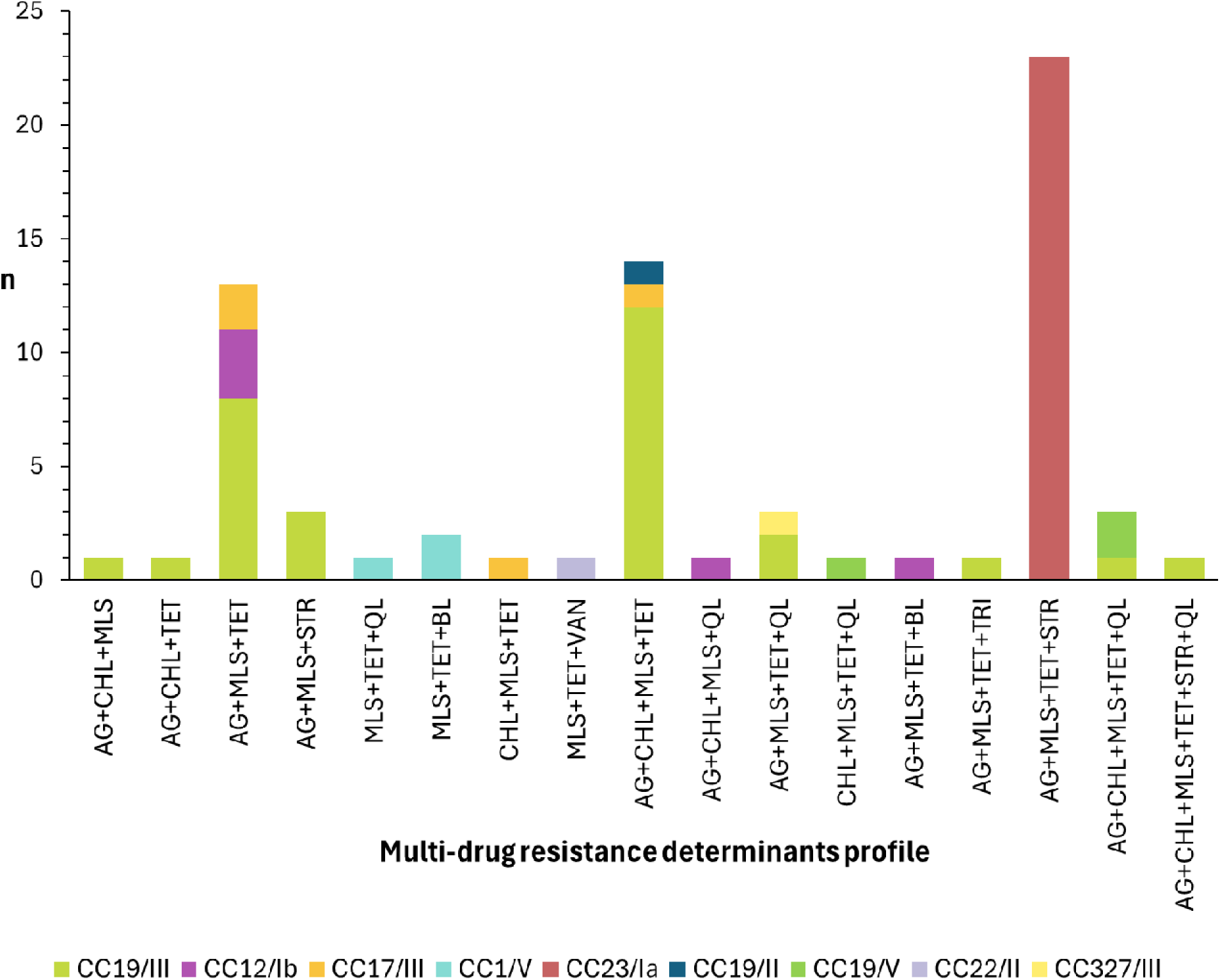
Distribution of GBS lineages according to multi-drug resistance determinants profile. AG: aminoglycosides; CHL: chloramphenicol; MLS: macrolides/lincosamides/streptogramins; TET: tetracyclines; STR: streptothricin; QL: quinolones; BL: beta-lactams; VAN: vancomycin; TRI: trimethoprim.

It is notable that ARDs to MLS were found in 70/71 isolates carrying ARDs to three or more antibiotic classes. Furthermore, of the 131 isolates with ARDs to MLS, 124 carried ARDs to at least one more antibiotic class, with ARDs to tetracyclines present in 116/124 of those isolates.

Among Argentinian isolates, uGBS carried a significantly (*p*<0.01) higher proportion of ARDs to multiple antibiotic classes than the rest of collections (41% vs 18-30%, Supplementary Figure 7). A single isolate carried ARDs to 6 classes and represented CC19/III niGBS.

## Discussion

There are few studies describing the epidemiology of GBS isolated from humans in South America, and even fewer that report serotype or clonal complex distribution. The most recent data is from Brazil, where serotype Ia was reported as the most prevalent, followed by serotypes II or Ib, in maternal colonisation, infant disease, and invasive and non-invasive infections in non-pregnant adults (70–74). All Argentinian GBS collections analysed in this study presented CC23/Ia and CC12/Ib as the dominant lineages. Lineage prevalence distribution in South America contrasts with reports from other parts of the world, where there are different dominant lineages according to sample source. For example, the most prevalent lineage associated with infant invasive infections worldwide, except for South America, is CC17/III, while for infections in adults the most prevalent lineage tends to be CC1/V (75–77). In the case of maternal colonisation, there is more heterogeneity in the distribution of the dominant lineages according to the region of the world (78–80). For instance, in Europe and China the predominant lineage is CC19/III (81,82), in most African regions it is CC1/V (83), while in Australia it is CC23/Ia (84). In spite of the difference in lineage distribution, Argentinian isolates were found to be related to GBS from around the globe and to belong to both host-adapted and host-generalist lineages, as many of the prevalent lineages, specifically CC23/Ia, CC12/Ib and CC1/V, have been previously detected in animal species in different countries (85–87).

Surveillance of serotype distribution in each region is key to development of capsule polysaccharide-based GBS vaccines to estimate their possible coverage. In this regard, phenotypically non-typable (pNT) GBS isolates are a cause of concern, as the coverage of the serotype-specific vaccines is uncertain. In different parts of the world pNT GBS were reported in a range of 5-17% (17,19,88–93), while in Argentina only 3% (12/365) of isolates were pNT. In 1/12 of the Argentinian pNT isolates, only 2 genes of the *cps* operon were found, indicating a loss of the capsule (90,94). In another of the pNT isolates, all genes of the capsular type III were found, but *cps*J was shorter, which could have led to low or lack of capsule expression (90). In the remaining 10/12 pNT isolates, a capsular type could be assigned by genomic analysis, which could indicate sporadic errors in the phenotypic methods or a low or lack of expression of the capsular genes (95). Discrepancies between the capsular genotype and the phenotypic expression or errors in the interpretation of agglutination results can also be found in phenotypically typeable GBS, as was reported in 10-29% of isolates from different parts of the world (84,96,97). In Argentinian isolates, discrepancy of 15% was detected with no correlation between the capsular type and the differing serotype, so we believe that they might be due to random events and not something intrinsic to the *cps* loci sequence or its expression in one particular capsular type, nor systematic errors in the phenotypic detection method.

Two vaccines against GBS are currently in the final stages of clinical development. Pfizer is working on a hexavalent conjugate vaccine (GBS6) containing the capsular polysaccharides of serotypes Ia, Ib, II, III, IV and V (98). As this vaccine is aimed at pregnant women, it would cover an estimated 99% of maternal colonisation cases in Argentina and it could prevent 99% of infant infections. Applied to an adult general population, it could also help prevent 96% of invasive infections and 99% of urinary infections caused by GBS. Following a different approach, MinervaX has designed a recombinant protein vaccine (GBS-NN/NN2) based on the N-terminal domains of the prevalent surface proteins of the Alp family (AlpCN, RibN, Alp1N and Alp2/3N), thus avoiding concerns related to variations in GBS capsular type (99). The genes coding for one or more of these proteins were found in over 97% of Argentinian isolates, so a very high coverage would be expected also from this vaccine.

Over 71% of Argentinian isolates carried at least one prophage, which is consistent with previous reports on prophage content in GBS in different parts of the world (32,33,35,100,101). No significant differences were found in the total prophage prevalence based on collection type, though a lower prophage content was observed in GBS recovered from infant invasive infections. This lack of significant difference might be attributed to the low number of iiGBS recovered, a likely positive outcome of GBS prenatal screening being mandatory in Argentina. These results contrast with a study of GBS collections from France, where it was reported that isolates from disease carried more prophages than colonising isolates (32,102). This could be explained by differences in the prevalent GBS lineages in France and Argentina, since prophage content is associated with GBS lineage, and we found no significant differences in the lineage distribution among isolates from disease and colonisation.

While no virulence or antibiotic resistance determinants commonly found in GBS were carried by the prophages analysed in this study, these prophages do carry genes involved in bacterial fitness, host adaptation to stressful environments and virulence, and up to 60% of ORFs encoding proteins with unknown function, as reported in our previous study (36). Interestingly, the highest density of prophages, virulence and antibiotic resistance determinants was found in the CC19 isolates, mostly of capsular type III, independently from the origin of the isolates (disease or colonisation).

The prophages of type A 12/111phiA and phiStag1 had been reported to be associated with the clonal expansion of CC17 isolates causing severe infant infections in France (33,61) and the Netherlands (34), respectively. However, no evidence of clonal expansion related to the presence of these prophages were found in Argentinian isolates. Most GBS isolates carrying these prophages belonged to CC23/Ia, but they shared the same phylogenetic subclades as those without A prophages.

The presence of prophages with integrase type GBS*Int*8.1 or GBS*Int*8.2 in isolates recovered from disease but not from carriage, does not seem to be associated with a higher invasiveness of their host lineage, as they were found in a wide variety of GBS lineages. Instead, it could be linked to the carriage by all the prophages (15/15) of a gene with a Phox Homology (PX) domain (36), which is a phosphoinositide binding domain involved in protein-membrane and protein-protein interactions in eukaryotic cells (103,104). The PX domain was not found in any of the prophages carried by colonisation isolates, so it is possible that prophages encoding a protein with this domain give the host bacteria an advantage to invade the human body. GBS*Int*8 prophages also have in common their insertion between a transcriptional regulatory protein (YbaB/EbfC family) and a hypothetical protein (36,50). It remains to be investigated whether their insertion site might alter the expression of genes that are beneficial for host invasion. In any case, more isolates from disease and carriage need to be analysed to confirm that these prophages are exclusive to disease-causing isolates.

Phenotypic antibiotic resistance rates in our GBS collections were previously discussed (3,5,59). In more than 97% of our isolates, the presence of ARDs correlated with the phenotypic resistance determined for MLS, aminoglycosides and quinolones. This shows the high predictability potential for antibiotic resistance from whole genome sequence analyses. In the case of the mutations in *pbp*2x found in this study, their presence was not associated with RBLS, but it cannot be ruled out that they could be a risk factor for the generation of a PBP2x with lower affinity for these antibiotics in the event of a new mutation or selection pressure (69).

The frequency of isolates with multiple ARDs is worrisome, especially those with combined resistance to MLS, aminoglycosides and tetracyclines (almost 13% of the Argentinian isolates). The combination of tetracycline and macrolides resistance determinants has already been described two decades ago, and it was proposed that tetracycline-resistant isolates could play a role in the dissemination of macrolide-resistant strains, as both types of genes are usually found in the same mobile elements (63). It needs to be further investigated if the aminoglycoside ARDs in our isolates are co-carried on the same genetic elements, as described in *Streptococcus pyogenes* (105). High prevalence of multidrug-resistant CC19/III GBS has been previously described, mainly associated with fluoroquinolone-resistance, and presence of mobile genetic elements (MEGs) carrying multiple resistance determinants were reported (106–111). Further analysis of MEGs present in the Argentinian GBS genomes is warranted to investigate co-carriage of ARDs to multiple antibiotics, which might promote transmission of multidrug resistance in our population.

GBS isolated from urinary tract infections (UTI) had the highest frequency of resistance determinants to aminoglycosides and novel variants of *pbp*2x. Isolates containing ARDs to quinolones but susceptible to LEV and NOR were also found in this collection. These findings suggest that GBS colonising the urinary tract is subjected to increased selective pressure due to the exposure to beta-lactams, fluoroquinolones and aminoglycosides excreted in the urine (112–114) following their use for the treatment of different kinds of infections. This heightened selective pressure, combined with the accumulation of mutations associated with antibiotic resistance, could increase the risk of clinical treatment failure for UTIs caused by GBS.

## Conclusions

This is the first analysis of human-isolated GBS population based on whole genome sequence data in South America, with special focus on the analysis of prophage content. The findings in this study suggest a possible association between an increased GBS virulence and the carriage of prophages with integrase type GBS*Int*8 and/or the presence of genes that encode the Phox Homology domain. Given the high prevalence of the serotypes and relevant virulence factors that are targeted by the in-development GBS vaccines, we conclude that both would be highly effective in preventing infections in all age-groups affected by GBS in Argentina. Implementation of prevention strategies against GBS infections is crucial, especially in the context of the rising GBS resistance to multiple antibiotic classes. Given the lack of genomic epidemiology data from human-isolated GBS in South America, this study makes a significant contribution to our understanding of the global GBS population structure.

## Ethical approval

Ethical approval for the ‘‘Argentinian multicentric study on infections due to Streptococcus agalactiae’’ was provided by the Ethics Committee of the Faculty of Pharmacy and Biochemistry, University of Buenos Aires, Res [D] RESCD-2022-400-E-UBA-DCT.

## Author Contributions

VK, SDG, MM and LB conceived this study and LB supervised it. TP, JC, UBK, DJ and SB conducted and coordinated the whole genome sequencing. VK and SDG designed the bioinformatic analyses. VK, MP and SDG performed the acquisition of the data. VK and SDG conducted the data analysis. SDG guided and supervised the bioinformatic analysis. CC participated in the discussion of the results and gave critical insights for their analysis. VK wrote the original draft of the manuscript. All authors reviewed, edited and approved the final version of the manuscript.

## Acknowledgments

We would like to thank all 40 participating centers of the multicentric study, their microbiologists and infectious disease specialists, and everyone involved in the genome sequencing of our GBS collection at the sequence facilities of the Wellcome Sanger Institute and Instituto Malbrán.

